## Supplemental Figures and Table for "The AAA+ chaperone VCP disaggregates Tau fibrils and generates aggregate seeds"

Supplementary information includes 10 Supplementary Figures and 1 Table

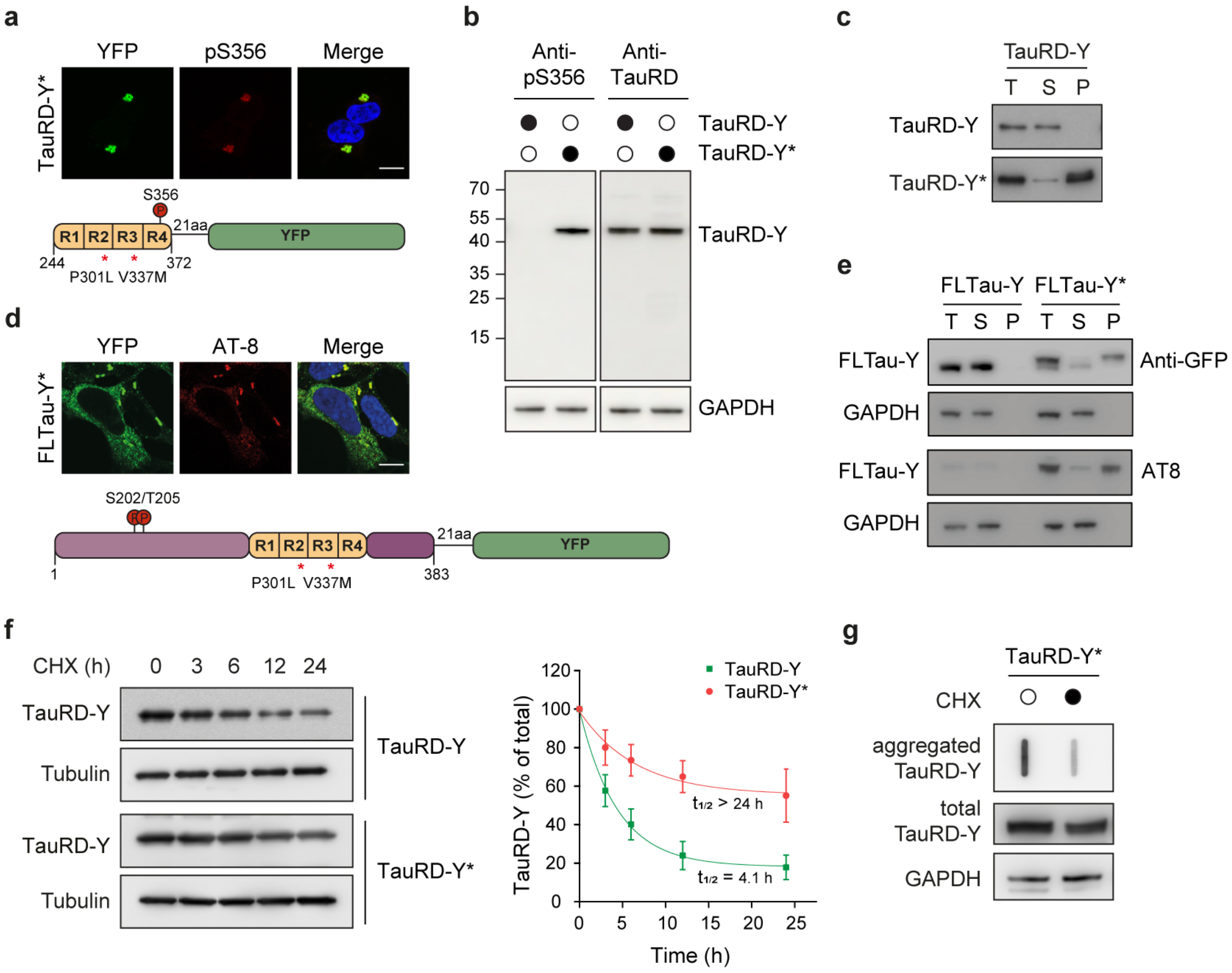


**Supplementary Fig. 1: Tau aggregation and clearance in a constitutive expression model.**
**a** Immunofluorescence staining of TauRD-Y* cells with an antibody against Tau phosphorylation at S356 (red) and YFP fluorescence of TauRD-Y (green). Scale bar, 10 µm. **b** Analysis of Tau S356 phosphorylation in lysates of TauRD-Y and TauRD-Y* cells by immunoblotting. Total TauRD-Y was detected using antibody against TauRD. **c** Solubility of TauRD-Y in TauRD-Y and TauRD-Y* cells at steady state, determined by fractionation of cell lysate by centrifugation, followed by immunoblotting with anti-GFP antibody. T, total cell lysate, S, supernatant, P, pellet. **d** Immunofluorescence staining of full-length Tau (FLTau-Y) in aggregate-containing FLTau-Y* cells with AT-8 antibody specific for Tau phosphorylation at S202/T205 (red) and YFP fluorescence of TauRD-Y (green). Scale bar, 10 µm. **e** Solubility of phosphorylated FLTau-Y in FLTau-Y and FLTau-Y* cells at steady state analyzed as in (c). Immunoblotting was with AT-8 antibody (bottom) and anti-GFP (top). GAPDH served as loading control. **f** Turnover of TauRD-Y in TauRD-Y and TauRD-Y* cells upon cycloheximide (CHX) shut-off (CHX; 50 µg/mL). Left, anti-GFP immunoblots to determine TauRD-Y levels. Tubulin served as loading control. Right, exponential fits of CHX chase data and corresponding half-lives ($t_{1/2}$). Mean ± s.d.; n=3. **g** Filter trap analysis of aggregated TauRD-Y upon CHX chase for 24 h. Aggregated and total TauRD-Y levels were determined by anti-GFP immunoblotting. GAPDH served as loading control.

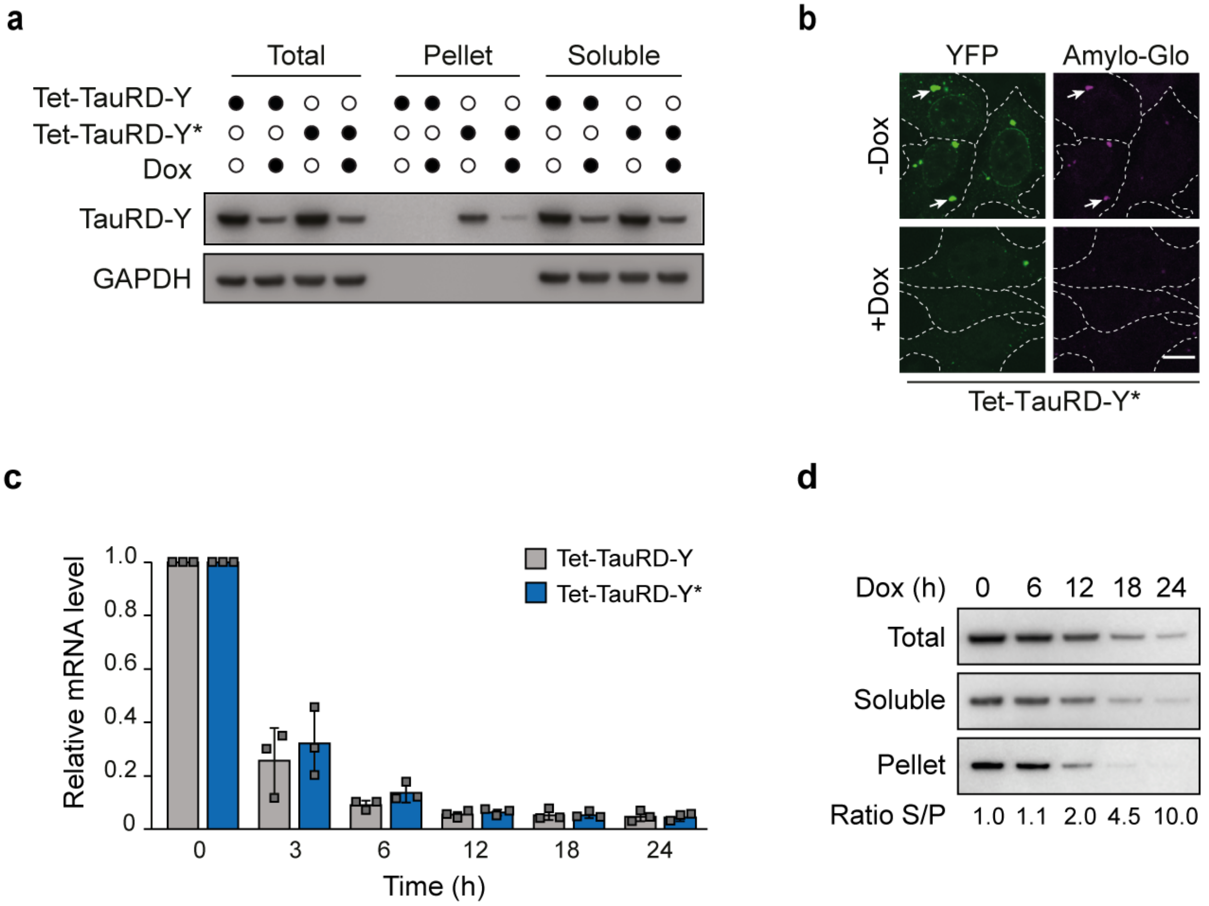


**Supplementary Fig. 2: TauRD-Y aggregation and clearance upon inhibition of expression in a Tet-regulated TauRD-Y expression system.**

**a** Solubility of TauRD-Y in Tet-TauRD-Y and Tet-TauRD-Y* cells upon addition of 50 ng/mL doxycycline (Dox) for 24 h. Cell lysates were fractionated as in Supplementary Fig. 1c. TauRD-Y was detected with anti-GFP antibody. GAPDH served as loading control. **b** Representative fluorescence images of Tet-TauRD-Y* cells treated with Dox for 24 h showing staining of TauRD-Y inclusions (green) with Amylo-Glo (magenta). White dashed lines indicate cell boundaries. Scale bar, 10 µm. **c** Quantitative PCR analysis of TauRD-Y mRNA in Tet-TauRD-Y and Tet-TauRD-Y* cells treated with Dox for 0, 3, 6, 12, 18 and 24 h. mRNA levels were normalized to the reference gene *RPS18*. Mean ± s.d.; n=3. **d** Solubility of TauRD-Y in Tet-TauRD-Y* cells upon addition of Dox for the indicated times. Normalized ratios of TauRD-Y in soluble (S) and pellet (P) fractions are stated.

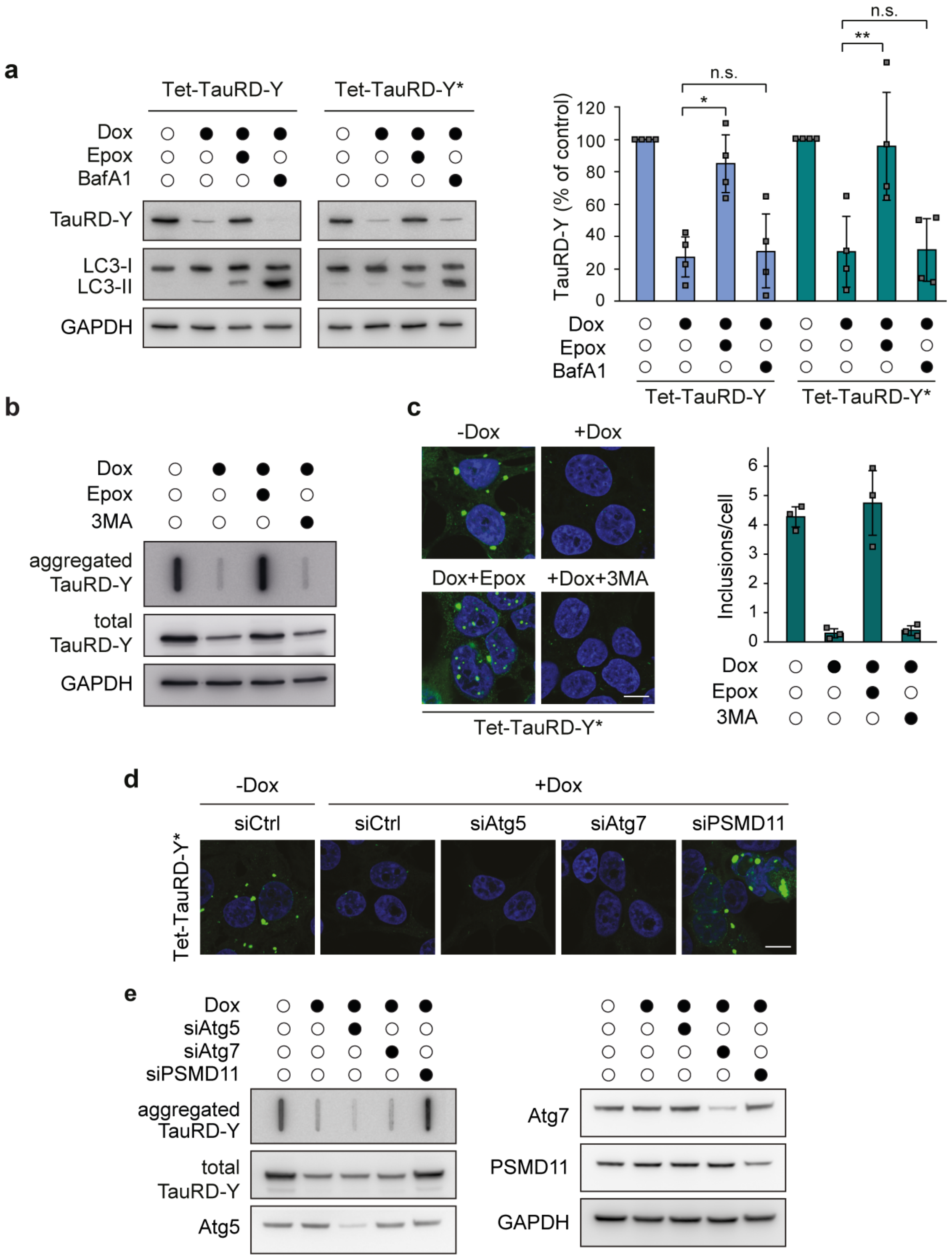


**Supplementary Fig. 3: Effect of UPS and autophagy inhibition on TauRD-Y levels and aggregate clearance.**

**a** Analysis of TauRD-Y levels in Tet-TauRD-Y and Tet-TauRD-Y* cells treated for 24 h with doxycycline (Dox; 50 ng/mL) alone or in combination with Epoxomicin (Epox; 50 nM) or

Bafilomycin A1 (BafA1; 50 nM). TauRD-Y and LC3 levels were determined by immunoblotting against GFP and LC3B respectively. GAPDH served as loading control. Mean ± s.d.; n=4. *p<0.05 (Tet-TauRD-Y: + Dox vs + Dox + Epox, p=0.0114), **p<0.01 (Tet-TauRD-Y*: + Dox vs + Dox + Epox, p=0.0026); n.s. non-significant (Tet-TauRD-Y: + Dox vs + Dox + Epox, p= 0.6422; Tet-TauRD-Y*: + Dox vs + Dox + Epox, p= 0.8799) from two-tailed Student's paired t-test. **b** Filter trap analysis of Tet-TauRD-Y* cells treated for 24 h with Dox alone or in combination with Epoxomicin (Epox; 50 nM) or 3-methyladenine (3MA; 5 mM). Aggregated and total TauRD-Y was detected with anti-GFP antibody. **c** Left, representative images of Tet-TauRD-Y* cells treated for 24 h with Dox alone or, in combination with Epoxomicin (Epox; 50 nM) or 3MA (5 mM). Scale bar, 10 µm. Right, quantification of TauRD-Y foci. 100-200 cells analyzed per experiment. Mean ± s.d.; n=3. **d** Representative images of Tet-TauRD-Y* cells transfected with non-targeted (Ctrl) siRNA or siRNA against Atg5 (50 nM), Atg7 (50 nM) and PSMD11 (25 nM). 72 h after transfection, doxycycline (Dox; 50 ng/mL) was added for another 24 h where indicated. Scale bar, 10 µm. **e** Filter trap analysis of Tet-TauRD-Y* cells transfected with siRNAs and treated with Dox as stated in (d). TauRD-Y was detected by immunoblotting with anti-GFP antibody.

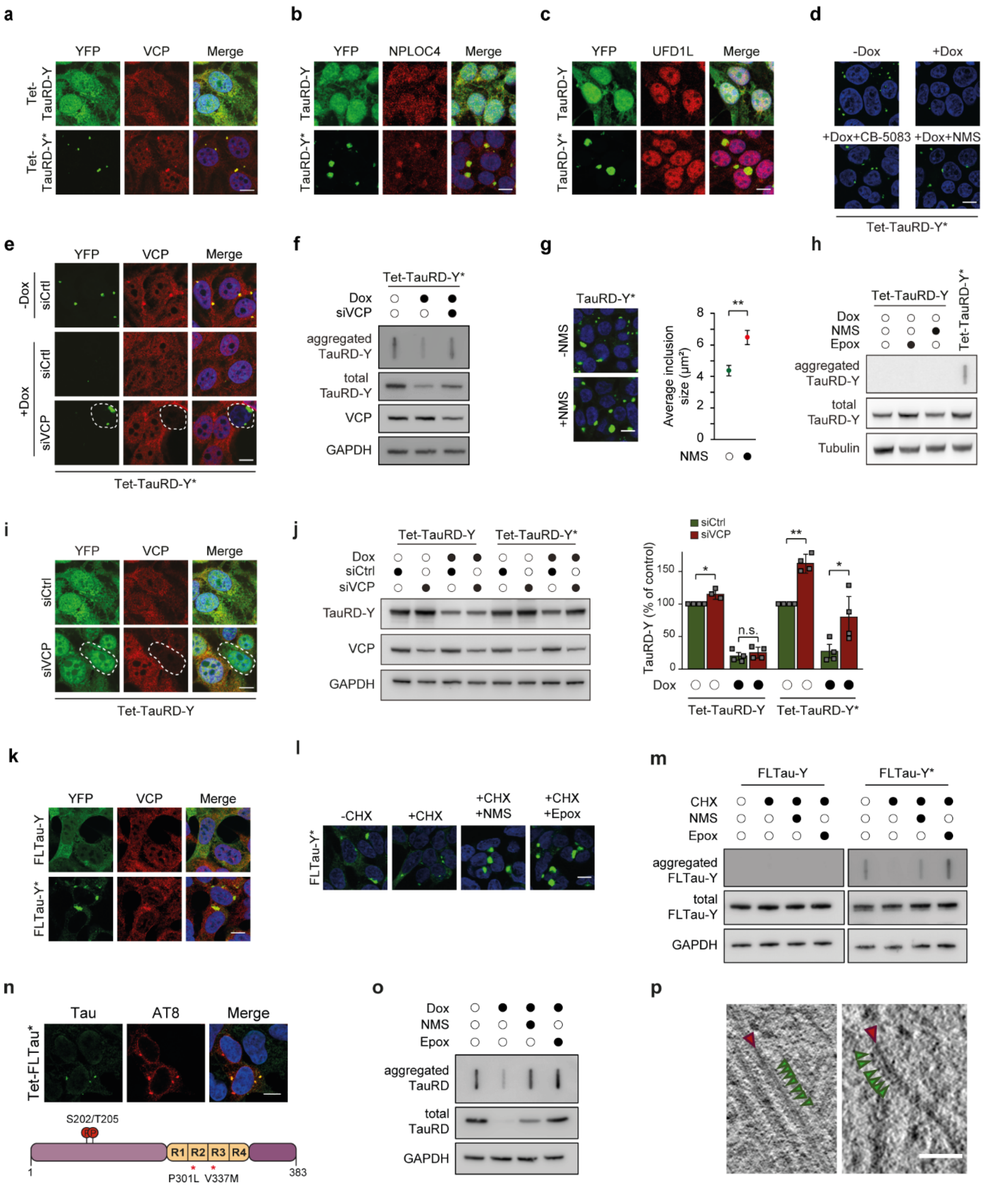


**Supplementary Fig. 4: Aggregation specific stabilization of Tau by VCP inactivation.**

**a** Immunofluorescence staining of VCP (red) and YFP fluorescence of TauRD-Y (green) in Tet-TauRD-Y and Tet-TauRD-Y* cells. **b** and **c** Immunofluorescence staining of NPLOC4 (b) (red) and UFD1L (c) (red) in TauRD-Y and TauRD-Y* cells. Scale bars, 10 µm. **d** Representative images of Tet-TauRD-Y* cells treated for 24 h with doxycycline (Dox; 50 ng/mL) alone or in combination with CB-5083 (1 μM) or NMS-873 (NMS; 2.5 μM). Scale bar, 10 μm. **e** Immunofluorescence staining of VCP (red) in Tet-TauRD-Y* cells treated with non-targeted (Ctrl) or VCP siRNA for 96 h. Doxycycline (Dox; 50 ng/mL) was added for the last 24 h. Dashed lines indicate a cell with reduced VCP levels. Scale bar, 10 μm. **f** Filter trap analysis of aggregated TauRD-Y in Tet-TauRD-Y* lysates treated as in (d). Aggregated and total TauRD-Y was analyzed by anti-GFP immunoblotting. GAPDH served as loading control. **g** Size increase of TauRD-Y inclusions upon VCP inhibition. Representative images of TauRD-Y* cells treated for 24 h with NMS-873 (NMS; 5 μM) and quantification of average inclusion size ($\mu m^2$). 200-400 cells analyzed per experiment. Mean ± s.d.; n=5. **p<0.01 (p= 0.0022) from two-tailed Student's paired t-test. **h** Filter trap analysis of Tet-TauRD-Y cells treated for 24 h with Epoxomicin (Epox; 50 nM) or NMS-873 (NMS; 2.5 μM) where indicated. Tet-TauRD-Y* lysate was used as control. TauRD-Y was detected by immunoblotting with anti-GFP antibody. **i** Immunofluorescence staining of VCP (red) and YFP fluorescence of TauRD-Y (green) in Tet-TauRD-Y cells transfected with non-targeted (Ctrl) or VCP siRNA for 96 h. Dashed lines indicate a cell with reduced VCP levels. Scale bar, 10 μm. **j** Left, analysis of TauRD-Y level in Tet-TauRD-Y and Tet-TauRD-Y* cells transfected for 96 h with non-targeted (Ctrl) or VCP siRNA where indicated. Doxycycline (Dox; 50 ng/mL) was added for the last 24 h. TauRD-Y was detected by immunoblotting with anti-GFP antibody. Right, quantification of TauRD-Y immunoblot. Mean ± s.d.; n=4. *p<0.05 (Tet-TauRD-Y - Dox: siCtrl vs siVCP, p= 0.0218; Tet-TauRD-Y* + Dox: siCtrl vs siVCP, p= 0.0156); **p<0.01 (Tet-TauRD-Y* - Dox: siCtrl vs siVCP, p= 0.0023); n.s. non-significant (Tet-TauRD-Y + Dox: siCtrl vs siVCP, p= 0.0539) from two-tailed paired Student's t-test. **k** Immunofluorescence staining of VCP (red) and YFP fluorescence of FLTau-Y (green) in FLTau-Y and FLTau-Y* cells. Scale bar, 10 μm. **l** Representative images of FLTau-Y* cells treated for 24 h with cycloheximide (CHX; 50 μg/mL) alone or in combination with NMS-873 (NMS; 2.5 μM) or Epoxomicin (Epox; 100 nM). Scale bar, 10 μm. **m** Filter trap analysis of lysates from FLTau-Y and FLTau-Y* cells treated for 24 h with Dox alone or in combination with NMS-873 (NMS; 2.5 μM) or Epoxomicin (Epox; 50 nM). Aggregated and total FLTau-Y levels were determined by immunoblotting against GFP. GAPDH served as loading control. **n** Immunofluorescence staining of full-length Tau (FLTau) in aggregate-containing Tet-FLTau* cells with Tau (green) and Tau S202/T205 phosphorylation specific AT-8 (red) antibody. Scale bar, 10 μm. **o** Filter trap analysis of lysates from Tet-TauRD* cells treated for 24 h with Dox alone or in combination with NMS-873 (NMS; 2.5 μM) or Epoxomicin (Epox; 50 nM). Aggregated and total TauRD levels were determined by immunoblotting against myc and TauRD, respectively. GAPDH served as loading control. **p** Examples of two TauRD-Y fibrils from a representative 1.4 nm thick tomographic slice of a TauRD inclusion from neurons. Red arrows indicate TauRD-Y fibrils and green arrows indicate globular densities along fibrils. Scale bar, 40 nm.

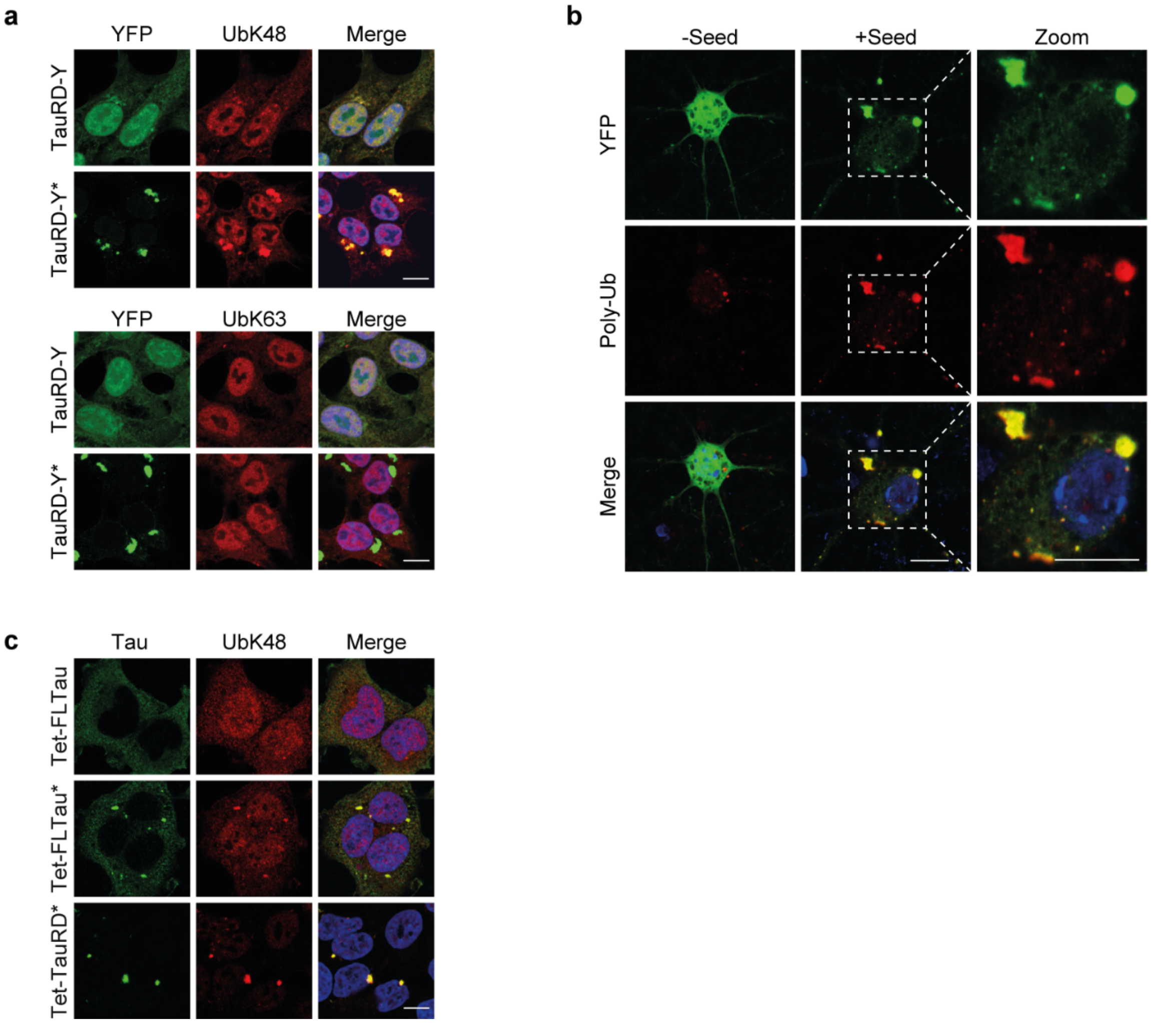


**Supplementary Fig. 5: Ubiquitylation of TauRD-Y aggregates.**

**a** Immunofluorescence staining of (top) ubiquitin-K48 (UbK48) (red) and (bottom) ubiquitin K63 (UbK63) (red) chains and YFP fluorescence of TauRD-Y (green) in TauRD-Y and TauRD-Y* cells. Scale bars, 10 µm. **b** Immunofluorescence staining of ubiquitylated proteins (FK2 antibody) (red) in primary neurons expressing TauRD-Y (green) and treated with TauRD containing lysates (+Seed) where indicated. Scale bars, 20 µm. **c** Immunofluorescence staining of ubiquitin-K48 (UbK48) (red) chains and Tau (green) in Tet-FLTau, Tet-FLTau* and Tet-TauRD* cells. FLTau was detected using Tau-5 and TauRD using anti-myc antibody. Scale bar, 10 µm.

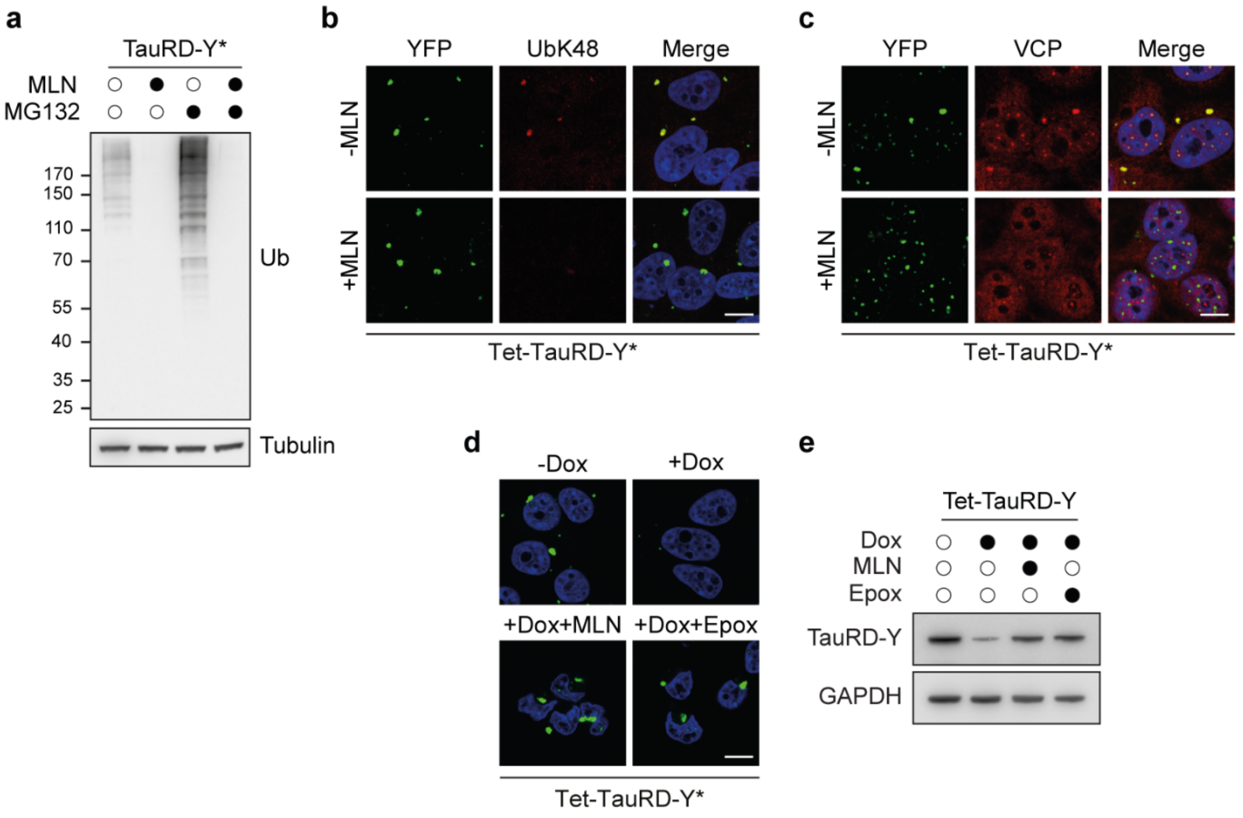


**Supplementary Fig. 6: Role of ubiquitylation in TauRD-Y disaggregation.**

**a** Analysis of ubiquitylated protein levels in lysates of TauRD-Y* cells treated with the ubiquitin activating enzyme E1 inhibitor MLN7243 (MLN; 0.5 µM) alone or in combination with proteasome inhibitor MG132 (1 µM) for 14 h. Ubiquitylated proteins were detected by immunoblotting against ubiquitin. Tubulin served as loading control. **b** Immunofluorescence staining of ubiquitin-K48 chains (UbK48) (red) and **c** VCP (red) in Tet-TauRD-Y* cells treated with MLN7243 (MLN; 0.5 µM) for 12 h. Scale bars, 10 µm. **d** Representative images of Tet-TauRD-Y* cells treated for 24 h with doxycycline (Dox; 50 ng/mL) alone or in combination with MLN7243 (MLN; 0.5 µM) or Epoxomicin (Epox; 50 nM). Scale bar, 10 µm. **e** Analysis of TauRD-Y levels in Tet-TauRD-Y cells treated for 24 h with Dox, MLN7243 and Epoxomicin as in (d). GAPDH served as loading control.

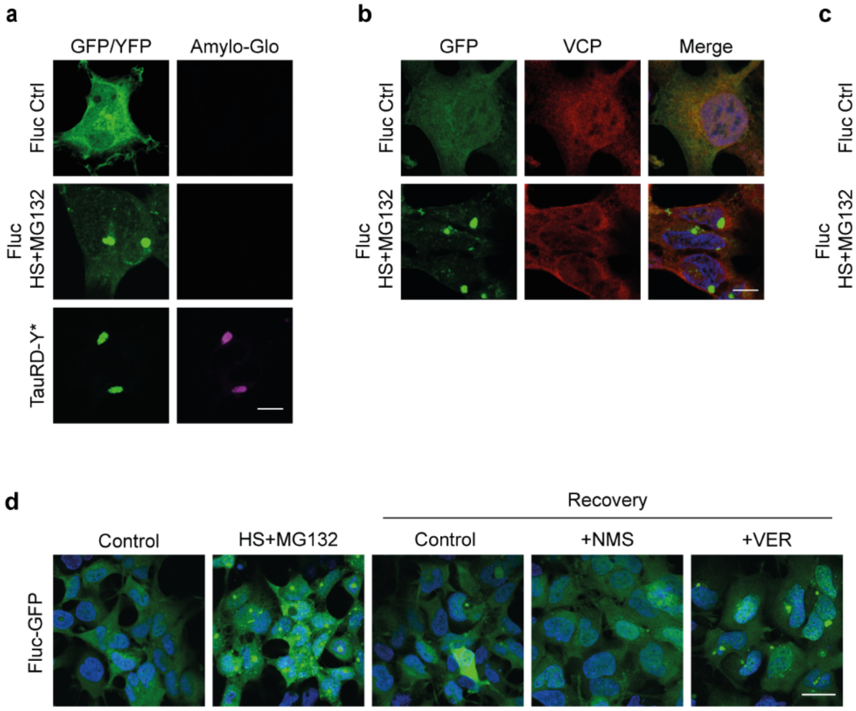


**Supplementary Fig. 7: Effect of VCP inhibition on firefly luciferase (Fluc) disaggregation.** **a** Fluc-GFP expressing cells maintained at 37 °C (Fluc Ctrl) or heat-stressed at 43 °C in presence of 5 µM MG132 for 2 h (Fluc HS) were stained with the amyloid-specific dye Amylo-Glo (magenta). TauRD-Y* cells were used as control. Amylo-Glo fluorescence was imaged with similar exposure settings in all panels. Scale bar, 10 µm. **b** Immunofluorescence staining of VCP (red), and **c** ubiquitylated proteins (FK2 antibody) (red) in Fluc-GFP cells treated as in (a). Scale bars, 10 µm. **d** Effect of VCP and Hsp70 inhibition on Fluc-GFP disaggregation. Fluc-GFP aggregation was induced as in (a). Cells were then shifted to MG132 free media and allowed to recover at 37 °C for 8 h in presence of NMS-873 (NMS; 2.5 µM) and VER-155008 (VER; 10 µM) where indicated. Scale bar, 30 µm.

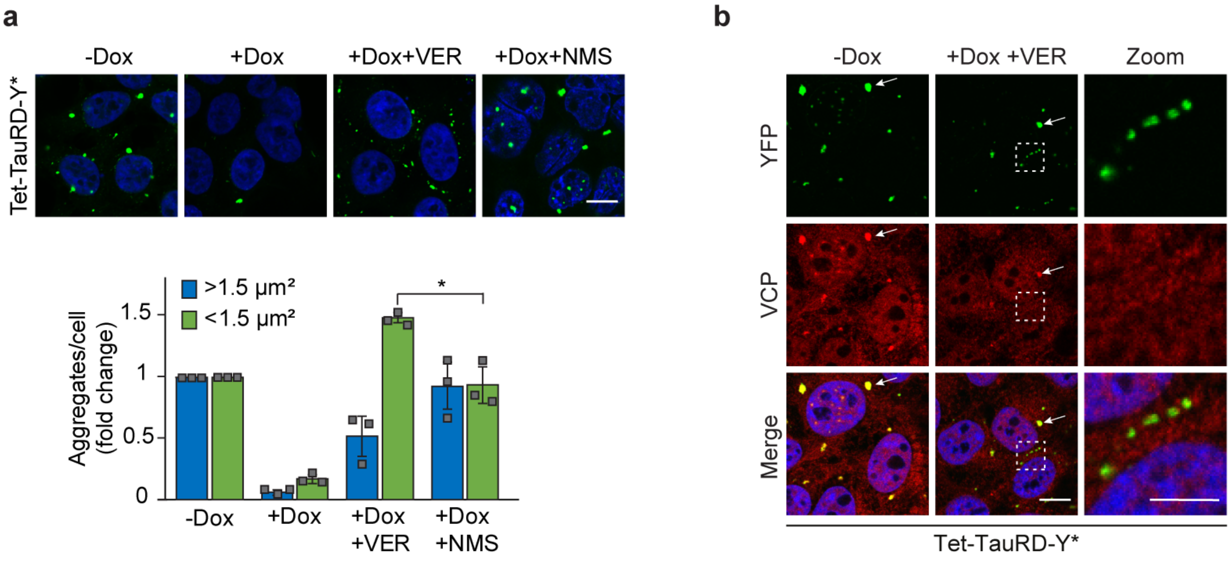


**Supplementary Fig. 8: Role of Hsp70 in TauRD-Y disaggregation.**

**a** Top, Representative images of Tet-TauRD-Y* cells treated for 24 h with doxycycline (Dox; 50 ng/mL) alone or in combination with VER-155008 (VER; 10 µM) or NMS-873 (NMS; 2.5 µM). Bottom, quantification of large (>1.5 µm$^2$) and small (<1.5 µm$^2$) TauRD-Y foci. Mean ± s.d.; n=3; ~100-200 cells were analyzed per experiment. *$p<0.05$ ($p=0.0435$) from two-tailed Student's paired t-test. Scale bar, 10 µm. **b** Immunofluorescence staining of VCP (red) and YFP fluorescence of TauRD-Y (green) in Tet-TauRD-Y* cells treated with a combination of doxycycline (Dox) and VER-155008 (VER) where indicated. White arrow points to large TauRD-Y inclusions co-localizing with VCP. Dashed lines enclose TauRD-Y foci that do not co-localize with VCP. Scale bar, 10 µm. Scale bar zoom, 5 µm.

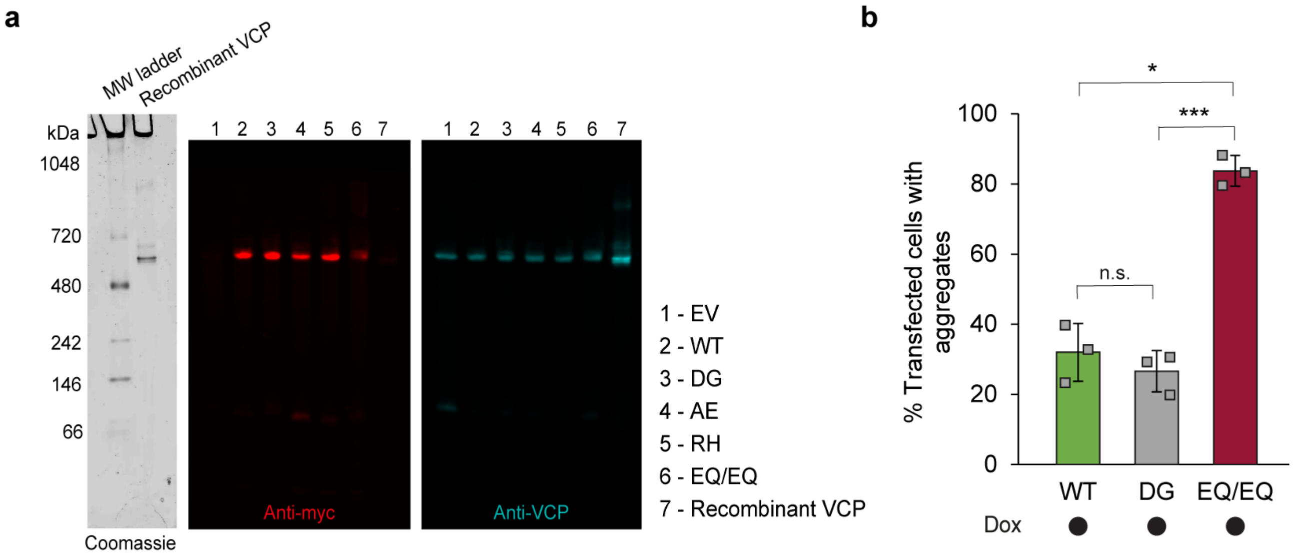


**Supplementary Fig. 9: Effect of VCP mutants on Tau disaggregation.**

**a** Native-PAGE analysis of recombinant VCP and lysates from Tet-TauRD-Y* cells transfected with empty vector (EV) and myc-tagged wild type (WT), D395G (DG), A232E (AE), R155H (RH) and E305Q/E578Q (EQ/EQ) VCP constructs. Immunoblot probed against myc (red) and VCP (cyan) is shown. Non-tagged, recombinant VCP was analyzed as control. **b** Quantification of aggregate foci in myc-positive Tet-TauRD-Y* cells transfected with myc-tagged WT, DG and EQ/EQ VCP constructs for 24 h, and treated for another 24 h with doxycycline (Dox; 50 ng/mL). Mean ± s.d.; n=3; > 100 cells analyzed per experiment; *$p<0.05$ (WT vs EQ/EQ $p=0.0192$); ***$p<0.001$ (DG vs EQ/EQ $p=0.0008$); n.s. non-significant ($p=0.5646$).

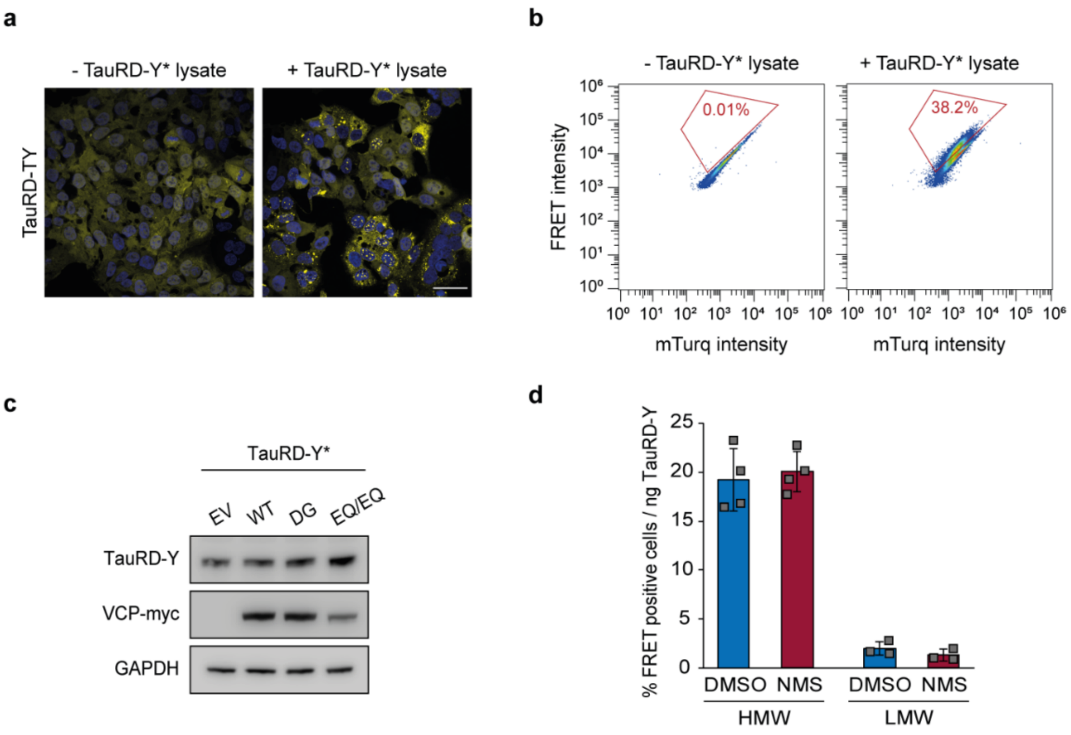


**Supplementary Fig. 10: Analysis of seeding-competent TauRD-Y.**

**a** Representative images of TauRD-TY FRET reporter cells treated with TauRD-Y* lysate where indicated showing TauRD-Y florescence in yellow. Scale bar, 40 µm. **b** Representative pseudocolour dot plots for the analysis of FRET positive TauRD-TY cells by flow cytometry upon addition of TauRD-Y* lysate. FRET intensity is plotted against mTurquoise2 (mTurq) intensity and the % of FRET positive cells are indicated in red gates. **c** Analysis of TauRD-Y and VCP-myc levels in TauRD-Y* cells transfected for two days with empty vector (EV) and myc-tagged wild type (WT), D395G (DG) and E305Q/E578Q (EQ/EQ) VCP constructs. TauRD-Y and overexpressed VCP levels were determined by immunoblotting against GFP and myc, respectively. GAPDH served as loading control. **d** Comparison of seeding efficiencies of high molecular weight (HMW) and low molecular weight (LMW) species obtained by size exclusion chromatography of lysates from TauRD-Y* cells treated for 24 h with DMSO or NMS-873 (NMS; 2 µM). Mean ± s.d.; HMW n=4, LMW n=3.

**Supplementary Table 1: TauRD-Y interactome in TauRD-Y* cells**

List of proteins interacting with TauRD-Y in TauRD-Y* cells at steady state determined by stable isotope labelling by amino acids in cell culture (SILAC). TauRD-Y cells were used as control. Normalized SILAC ratios of TauRD-Y*/TauRD-Y [H/L] from 3 independent replicates are shown. Interactors were defined as proteins quantified in at least 2 out of 3 replicates with enrichment ≥2 fold. Proteins with known association to the ubiquitin-proteasome system are highlighted in green and VCP-cofactor complex in red.

Intensity-based absolute quantification (iBAQ) values reflect measured molar protein amounts.

| Protein ID | Protein name | Gene name | Fasta headers | Unique peptides | Mol. weight [kDa] | Norm. ratio [H/L] Rep.1 | Norm. ratio [H/L] Rep.2 | Norm. ratio [H/L] Rep.3 | Median norm. ratio | iBAQ [H] Exp.1 | iBAQ [H] Exp.2 | iBAQ [H] Exp.3 |
|---|---|---|---|---|---|---|---|---|---|---|---|---|
| Q9UNZ2;F2Z2K | NSFL1 cofactor p47 | NSFL1C | sp\|Q9UNZ2\|NSF1C_HUMAN NSFL1 cofactor p47 OS=Homo sapi | 14 | 40,572 | 42,702 | 80,022 | 11,719 | 42,702 | 59143000 | 112180000 | 13783000 |
| Q9UQN3;A0A0 | Charged multivesicular body protein 2 | CHMP2B | sp\|Q9UQN3\|CHM2B_HUMAN Charged multivesicular body pro | 4 | 23,906 | 18,201 | 25,514 | 11,839 | 18,201 | 2093000 | 5417800 | 15472000 |
| P55072;C9IZA | Transitional endoplasmic reticulum AT | VCP | sp\|P55072\|TERA_HUMAN Transitional endoplasmic reticulum | 47 | 89,321 | 16,959 | 43,344 | 6,3181 | 16,959 | 29193000 | 94165000 | 26411000 |
| Q9UNM6;J3KN | 26S proteasome non-ATPase regulatory | PSMD13 | sp\|Q9UNM6\|PSD13_HUMAN 26S proteasome non-ATPase regu | 22 | 42,945 | 17,963 | 16,071 | 13,583 | 16,071 | 9313200 | 17290000 | 5085700 |
| Q6PEV8 | Protein FAM199X | FAM199X | sp\|Q6PEV8\|F199X_HUMAN Protein FAM199X OS=Homo sapien | 5 | 42,801 | 41,449 | 15,154 | 15,115 | 15,154 | 9838300 | 30031000 | 10267000 |
| F8VUA2;Q9HD | Charged multivesicular body protein 1 | CHMP1A | tr\|F8VUA2\|F8VUA2_HUMAN Charged multivesicular body prot | 5 | 19,531 | 14,648 | 19,186 | 12,293 | 14,648 | 25912000 | 113020000 | 38592000 |
| Q15773;F5H0Y | Myeloid leukemia factor 2 | MLF2 | sp\|Q15773\|MLF2_HUMAN Myeloid leukemia factor 2 OS=Hom | 6 | 28,147 | 16,867 | 13,819 | 6,564 | 13,819 | 74907000 | 153440000 | 42871000 |
| O43242;H0YG | 26S proteasome non-ATPase regulatory | PSMD3 | sp\|O43242\|PSMD3_HUMAN 26S proteasome non-ATPase regul | 34 | 60,977 | 13,413 | 15,879 | 6,9978 | 13,413 | 21734000 | 50167000 | 10257000 |
| O00231;J3QRY | 26S proteasome non-ATPase regulatory | PSMD11 | sp\|O00231\|PSD11_HUMAN 26S proteasome non-ATPase regul | 26 | 47,463 | 13,086 | 22,385 | 7,5615 | 13,086 | 14133000 | 35102000 | 7362100 |
| P62191;G3V4X | 26S protease regulatory subunit 4 | PSMC1 | sp\|P62191\|PRS4_HUMAN 26S proteasome regulatory subunit | 20 | 49,184 | 12,739 | 13,685 | 6,7346 | 12,739 | 6082800 | 18968000 | 4723500 |
| P17480 | Nucleolar transcription factor 1 | UBTF | sp\|P17480\|UBF1_HUMAN Nucleolar transcription factor 1 OS= | 22 | 89,405 | 12,209 | 11,585 | 7,1602 | 11,585 | 94071000 | 144030000 | 43389000 |
| O14545;F8VNX | TRAF-type zinc finger domain-containin | TRAFD1 | sp\|O14545\|TRAD1_HUMAN TRAF-type zinc finger domain-cont | 12 | 64,84 | 7,2319 | 19,2 | 11,522 | 11,522 | 6914300 | 16022000 | 16301000 |
| Q13501;E7EM | Sequestosome-1 | SQSTM1 | sp\|Q13501\|SQSTM_HUMAN Sequestosome-1 OS=Homo sapien | 16 | 47,687 | 9,6152 | 17,975 | 11,41 | 11,41 | 27933000 | 113830000 | 115270000 |
| Q13200;C9JPC | 26S proteasome non-ATPase regulatory | PSMD2 | sp\|Q13200\|PSMD2_HUMAN 26S proteasome non-ATPase regul | 46 | 100,2 | 7,2067 | 22,927 | 11,267 | 11,267 | 4555900 | 9942100 | 6380700 |
| H0Y6K2;A0A14 | Bromodomain-containing protein 2 | BRD2 | tr\|H0Y6K2\|H0Y6K2_HUMAN Bromodomain-containing protei | 11 | 88,288 | 24,922 | 7,6855 | 10,958 | 10,958 | 6685300 | 16169000 | 6613700 |
| H0YFD6;P4093 | Trifunctional enzyme subunit alpha, mi | HADHA | tr\|H0YFD6\|H0YFD6_HUMAN Trifunctional enzyme subunit alp | 29 | 86,371 | 10,902 | 11,422 | 8,175 | 10,902 | 45516000 | 92103000 | 31426000 |
| P55036;Q5VW | 26S proteasome non-ATPase regulatory | PSMD4 | sp\|P55036\|PSMD4_HUMAN 26S proteasome non-ATPase regul | 14 | 40,736 | 12,228 | 10,9 | 3,7986 | 10,9 | 19080000 | 82099000 | 8516700 |
| Q6QNY1;J3QRU | Biogenesis of lysosome-related organell | BLOC1S2 | sp\|Q6QNY1\|BL1S2_HUMAN Biogenesis of lysosome-related org | 3 | 15,961 | 10,78 | 15,848 | 2,0051 | 10,78 | 1892700 | 3652000 | 1718300 |
| P55084;F5GZQ | Trifunctional enzyme subunit beta, mit | HADHB | sp\|P55084\|ECHB_HUMAN Trifunctional enzyme subunit beta, | 25 | 51,294 | 9,9693 | 11,079 | 8,0079 | 9,9693 | 65103000 | 128900000 | 39724000 |
| P25788;G3V4X | Proteasome subunit alpha type-3 | PSMA3 | sp\|P25788\|PSA3_HUMAN Proteasome subunit alpha type-3 O | 9 | 28,433 | 3,0605 | 14,754 | 9,9671 | 9,9671 | 2846500 | 1743900 | 2866700 |
| A8MUA9;A8MU | Small ubiquitin-related modifier 4;Sma | SUMO3;SUMO4; | tr\|A8MUA9\|A8MUA9_HUMAN SMT3 suppressor of mif two 3 h | 1 | 15,317 | 8,4377 | 11,126 | | 9,78185 | 14432000 | 26087000 | 0 |
| Q8TAT6;I3L4U | Nuclear protein localization protein 4 | NPLOC4 | sp\|Q8TAT6\|NPL4_HUMAN Nuclear protein localization protein | 17 | 68,119 | 3,9291 | 15,451 | | 9,69005 | 1208400 | 5604000 | 0 |
| P25789;H0YM | Proteasome subunit alpha type-4;Prote | PSMA4 | sp\|P25789\|PSA4_HUMAN Proteasome subunit alpha type-4 O | 10 | 29,483 | 2,7706 | | 16,307 | 9,5388 | 4339400 | 0 | 4387000 |
| Q9UHD9 | Ubiquilin-2 | UBQLN2 | sp\|Q9UHD9\|UBQL2_HUMAN Ubiquilin-2 OS=Homo sapiens OX | 11 | 65,695 | 6,6843 | 11,794 | | 9,23915 | 8201000 | 24600000 | 108990 |
| O14818;H0Y58 | Proteasome subunit alpha type-7 | PSMA7 | sp\|O14818\|PSA7_HUMAN Proteasome subunit alpha type-7 O | 14 | 27,887 | 2,6045 | 9,1585 | 10,57 | 9,1585 | 3845500 | 10551000 | 10021000 |
| Q9UID3;E9PJ3 | Vacuolar protein sorting-associated pr | VPS51 | sp\|Q9UID3\|VPS51_HUMAN Vacuolar protein sorting-associate | 11 | 86,041 | 12,8 | 5,2875 | | 9,04375 | 603160 | 3513600 | 552200 |
| Q14596;B7Z5R | Next to BRCA1 gene 1 protein | NBR1 | sp\|Q14596\|NBR1_HUMAN Next to BRCA1 gene 1 protein OS=H | 12 | 107,41 | | 15,113 | 2,4528 | 8,7829 | 0 | 12693000 | 12934000 |
| Q92890;C9JNP | Ubiquitin fusion degradation protein 1 | UFD1L | sp\|Q92890\|UFD1_HUMAN Ubiquitin recognition factor in ER- | 9 | 34,5 | 11,411 | 6,134 | | 8,7725 | 785590 | 5835400 | 721170 |
| Q15008;C9J0E | 26S proteasome non-ATPase regulatory | PSMD6 | sp\|Q15008\|PSMD6_HUMAN 26S proteasome non-ATPase regul | 27 | 45,531 | 7,3639 | 9,6008 | 8,555 | 8,555 | 13494000 | 32338000 | 6986900 |
| Q96DX7 | Tripartite motif-containing protein 44 | TRIM44 | sp\|Q96DX7\|TRI44_HUMAN Tripartite motif-containing protei | 5 | 38,472 | 10,369 | 6,5983 | | 8,48365 | 708170 | 19187000 | 783500 |
| Q16643;D6R9 | Drebrin | DBN1 | sp\|Q16643\|DREB_HUMAN Drebrin OS=Homo sapiens OX=960 | 16 | 71,428 | | 13,93 | 2,7427 | 8,33635 | 123350 | 13631000 | 6342500 |
| A0A087X2I1;P | 26S protease regulatory subunit 10B | PSMC6 | tr\|A0A087X2I1\|A0A087X2I1_HUMAN 26S proteasome regulat | 16 | 45,796 | 10,533 | 6,0398 | | 8,2864 | 2266900 | 10434000 | 743130 |
| P51665;H3BNT | 26S proteasome non-ATPase regulatory | PSMD7 | sp\|P51665\|PSMD7_HUMAN 26S proteasome non-ATPase regul | 14 | 37,025 | 8,2466 | 13,713 | 6,6956 | 8,2466 | 9133800 | 34517000 | 6064400 |
| P35998;A0A1 | 26S protease regulatory subunit 7 | PSMC2 | sp\|P35998\|PRS7_HUMAN 26S proteasome regulatory subunit | 24 | 48,633 | 7,5017 | 8,0742 | 8,5178 | 8,0742 | 8531200 | 12549000 | 5021100 |
| B3KVL5;Q8TBK | Zinc finger CCHC domain-containing pr | ZCCHC10 | tr\|B3KVL5\|B3KVL5_HUMAN Zinc finger, CCHC domain contain | 1 | 20,308 | | 7,1775 | 8,4956 | 7,83655 | 18527000 | 60508000 | 11466000 |
| Q8IXW5 | Putative RNA polymerase II subunit B1 | RPAP2 | sp\|Q8IXW5\|RPAP2_HUMAN Putative RNA polymerase II subun | 13 | 69,508 | 10,475 | 7,8181 | 3,3996 | 7,8181 | 1205400 | 8566200 | 4524400 |
| P62979;J3QS3 | Ubiquitin-40S ribosomal protein S27a; | RPS27A;UBB;UB | sp\|P62979\|RS27A_HUMAN Ubiquitin-40S ribosomal protein S | 10 | 17,965 | 9,4683 | 7,6945 | 4,1911 | 7,6945 | 3164100000 | 7019100000 | 1951400000 |
| Q99460;A0A08 | 26S proteasome non-ATPase regulatory | PSMD1 | sp\|Q99460\|PSMD1_HUMAN 26S proteasome non-ATPase regul | 42 | 105,84 | 6,684 | 7,415 | 7,8358 | 7,415 | 3058500 | 12531000 | 2851500 |
| P43686 | 26S protease regulatory subunit 6B | PSMC4 | sp\|P43686\|PRS6B_HUMAN 26S proteasome regulatory subuni | 21 | 47,366 | 8,4576 | 2,5541 | 6,7907 | 6,7907 | 1137200 | 10458000 | 617140 |
| R4GMR5;K7EJR | 26S proteasome non-ATPase regulatory | PSMD8 | tr\|R4GMR5\|R4GMR5_HUMAN 26S proteasome non-ATPase reg | 10 | 32,551 | 1,3779 | 6,6145 | 13,37 | 6,6145 | 491820 | 3751000 | 6211100 |
| P0CAP2;H8Y6P | DNA-directed RNA polymerase II subuni | POLR2M;GCOM1 | sp\|P0CAP2\|GRL1A_HUMAN DNA-directed RNA polymerase II su | 4 | 41,739 | 6,9552 | 6,1579 | | 6,55655 | 2999600 | 6478700 | 0 |
| P49721;A0A08 | Proteasome subunit beta type-2 | PSMB2 | sp\|P49721\|PSB2_HUMAN Proteasome subunit beta type-2 OS= | 10 | 22,836 | 2,6138 | 10,075 | | 6,3444 | 526870 | 3115400 | 1949400 |
| Q9UQ35;I3L4D | Serine/arginine repetitive matrix protei | SRRM2 | sp\|Q9UQ35\|SRRM2_HUMAN Serine/arginine repetitive matrix | 44 | 299,61 | 8,8182 | 4,1061 | 6,3418 | 6,3418 | 40059000 | 69443000 | 34477000 |
| O75487 | Glypican-4;Secreted glypican-4 | GPC4 | sp\|O75487\|GPC4_HUMAN Glypican-4 OS=Homo sapiens OX=9 | 13 | 62,411 | 6,1176 | 9,697 | 5,2264 | 6,1176 | 12306000 | 31449000 | 4359700 |
| Q9H307;G3V5F | Pinin | PNN | sp\|Q9H307\|PININ_HUMAN Pinin OS=Homo sapiens OX=9606 | 39 | 81,627 | 6,4613 | 5,9829 | 3,8631 | 5,9829 | 180190000 | 359840000 | 205880000 |
| Q99615;K7ESP | DnaJ homolog subfamily C member 7 | DNAJC7 | sp\|Q99615\|DNJC7_HUMAN DnaJ homolog subfamily C membe | 35 | 56,44 | 32,017 | 5,9433 | 3,8506 | 5,9433 | 58381000 | 56886000 | 13723000 |
| O95816 | BAG family molecular chaperone regul | BAG2 | sp\|O95816\|BAG2_HUMAN BAG family molecular chaperone re | 9 | 23,772 | 5,7371 | 8,5666 | 3,8905 | 5,7371 | 10758000 | 36658000 | 8833600 |
| Q9BYN8 | 28S ribosomal protein S26, mitochond | MRPS26 | sp\|Q9BYN8\|RT26_HUMAN 28S ribosomal protein S26, mitoch | 7 | 24,211 | 5,6963 | 5,8954 | 3,1697 | 5,6963 | 15879000 | 55778000 | 11868000 |
| P10644;K7EPB | cAMP-dependent protein kinase type I- | PRKAR1A | sp\|P10644\|KAP0_HUMAN cAMP-dependent protein kinase typ | 12 | 42,981 | 5,6698 | 8,1797 | 2,4289 | 5,6698 | 1804900 | 8873600 | 3456600 |
| A0A0C4DG62;F | ADP-ribosylation factor-like protein 6-i | ARL6IP4 | tr\|A0A0C4DG62\|A0A0C4DG62_HUMAN ADP-ribosylation fact | 5 | 24,591 | 5,6534 | 5,8168 | 2,4357 | 5,6534 | 11791000 | 37307000 | 7853000 |
| Q15545 | Transcription initiation factor TFIID su | TAF7 | sp\|Q15545\|TAF7_HUMAN Transcription initiation factor TFIID | 5 | 40,259 | 6,7383 | 4,5475 | | 5,6429 | 1917000 | 3824800 | 262590 |
| F5H442;Q9981 | Tumor susceptibility gene 101 protein | TSG101 | tr\|F5H442\|F5H442_HUMAN Tumor susceptibility gene 101 pr | 8 | 40,917 | 6,9756 | 3,423 | | 5,1993 | 1298100 | 2512100 | 0 |

| | | | | | | | | | | | | |
|---|---|---|---|---|---|---|---|---|---|---|---|---|
| Q14677;H0YD5 | Clathrin interactor 1 | CLINT1 | sp\|Q14677\|EPN4_HUMAN Clathrin interactor 1 OS=Homo sapi | 17 | 68,259 | 4,5671 | 5,1051 | | 4,8361 | 483270 | 1390800 | 0 |
| Q16531;F5GY5 | DNA damage-binding protein 1 | DDB1 | sp\|Q16531\|DDB1_HUMAN DNA damage-binding protein 1 OS= | 51 | 126,97 | 5,1552 | 4,8124 | 4,7395 | 4,8124 | 2260500 | 2015000 | 818320 |
| O14646;H0Y8V | Chromodomain-helicase-DNA-binding | CHD1 | sp\|O14646\|CHD1_HUMAN Chromodomain-helicase-DNA-bind | 22 | 196,69 | 4,667 | 5,6909 | 3,5182 | 4,667 | 1212300 | 6836700 | 855680 |
| G3V5Z7;P6090 | Proteasome subunit alpha type;Proteas | PSMA6 | tr\|G3V5Z7\|G3V5Z7_HUMAN Proteasome subunit alpha type O | 12 | 28,147 | | 5,4361 | 3,7845 | 4,6103 | 0 | 986560 | 3211200 |
| P17980;E9PM6 | 26S protease regulatory subunit 6A | PSMC3 | sp\|P17980\|PRS6A_HUMAN 26S protease regulatory subuni | 21 | 49,203 | 5,2382 | 3,8803 | | 4,55925 | 2297800 | 7983700 | 563940 |
| O00232;J3KTJ5 | 26S proteasome non-ATPase regulatory | PSMD12 | sp\|O00232\|PSD12_HUMAN 26S proteasome non-ATPase regul | 22 | 52,904 | 4,4642 | 2,2919 | 13,215 | 4,4642 | 5157500 | 5534500 | 3641700 |
| P61964;V9GZ5 | WD repeat-containing protein 5 | WDR5 | sp\|P61964\|WDR5_HUMAN WD repeat-containing protein 5 OS | 5 | 36,588 | 2,7665 | 5,5202 | | 4,14335 | 1116200 | 6218500 | 163100 |
| O00487;C9JW3 | 26S proteasome non-ATPase regulatory | PSMD14 | sp\|O00487\|PSDE_HUMAN 26S proteasome non-ATPase regulat | 10 | 34,577 | 4,4008 | 3,6454 | | 4,0231 | 2763600 | 4175800 | 1273000 |
| P62195;J3QQ | 26S protease regulatory subunit 8 | PSMC5 | sp\|P62195\|PRS8_HUMAN 26S proteasome regulatory subunit | 23 | 45,626 | 4,5333 | 4,0178 | 2,4837 | 4,0178 | 2401900 | 10162000 | 3258800 |
| P24928 | DNA-directed RNA polymerase II subuni | POLR2A | sp\|P24928\|RPB1_HUMAN DNA-directed RNA polymerase II sub | 51 | 217,17 | 3,9675 | 4,1625 | 3,3865 | 3,9675 | 5036400 | 22719000 | 4400000 |
| O14974;F8VZN | Protein phosphatase 1 regulatory subu | PPP1R12A | sp\|O14974\|MYPT1_HUMAN Protein phosphatase 1 regulatory | 15 | 115,28 | 3,6323 | 4,1458 | | 3,88905 | 135210 | 738040 | 0 |
| Q5VIR6;F6VX9 | Vacuolar protein sorting-associated pr | VPS53 | sp\|Q5VIR6\|VPS53_HUMAN Vacuolar protein sorting-associated | 10 | 79,652 | | 2,3872 | 5,2434 | 3,8153 | 0 | 867750 | 158630 |
| Q8WV44;H0Y9 | E3 ubiquitin-protein ligase TRIM41 | TRIM41 | sp\|Q8WV44\|TRI41_HUMAN E3 ubiquitin-protein ligase TRIM4 | 5 | 71,669 | 4,3208 | 3,4885 | 3,7734 | 3,7734 | 613000 | 252640 | 1261800 |
| F8W118;F8VV59;B7Z9C2;F8VRJ2 | | NAP1L1 | tr\|F8W118\|F8W118_HUMAN Nucleosome assembly protein 1- | 12 | 24,694 | 3,6447 | 4,8952 | 3,4073 | 3,6447 | 4807500 | 20158000 | 2227100 |
| E9PNW4;A0A2 | CD59 glycoprotein | CD59 | tr\|E9PNW4\|E9PNW4_HUMAN Uncharacterized protein OS=Ho | 4 | 11,985 | | 4,7462 | 2,1357 | 3,44095 | 5131200 | 54954000 | 35045000 |
| Q96BQ5 | Coiled-coil domain-containing protein | CCDC127 | sp\|Q96BQ5\|CC127_HUMAN Coiled-coil domain-containing pro | 4 | 30,834 | 4,0401 | | 2,7807 | 3,4104 | 3768100 | 3178900 | 697410 |
| O43164 | E3 ubiquitin-protein ligase Praja-2 | PJA2 | sp\|O43164\|PJA2_HUMAN E3 ubiquitin-protein ligase Praja-2 C | 15 | 78,213 | 3,3397 | 6,1296 | 2,2392 | 3,3397 | 4791500 | 29710000 | 8235700 |
| B8ZZD4;Q86VP | Tax1-binding protein 1 | TAX1BP1 | tr\|B8ZZD4\|B8ZZD4_HUMAN Tax1-binding protein 1 OS=Homo | 14 | 93,609 | 9,5213 | 3,2831 | 3,1608 | 3,2831 | 409770 | 4896800 | 4112000 |
| P62857 | 40S ribosomal protein S28 | RPS28 | sp\|P62857\|RS28_HUMAN 40S ribosomal protein S28 OS=Hom | 2 | 7,8409 | 3,7217 | 3,2677 | 1,1842 | 3,2677 | 18987000 | 32783000 | 6557100 |
| P62873;F6UT2 | Guanine nucleotide-binding protein G( | GNB1 | sp\|P62873\|GBB1_HUMAN Guanine nucleotide-binding protei | 13 | 37,377 | 3,2364 | 4,2746 | 1,6165 | 3,2364 | 2812200 | 17175000 | 35389000 |
| Q7L7X3;J3QS7 | Serine/threonine-protein kinase TAO1 | TAOK1 | sp\|Q7L7X3\|TAOK1_HUMAN Serine/threonine-protein kinase T | 15 | 116,07 | 1,3856 | 3,2072 | 3,0804 | 3,0804 | 2889600 | 6633200 | 6110200 |
| O75955;A0A14 | Flotillin-1 | FLOT1 | sp\|O75955\|FLOT1_HUMAN Flotillin-1 OS=Homo sapiens OX=96 | 16 | 47,355 | 3,0802 | 3,6951 | 1,7534 | 3,0802 | 11366000 | 21730000 | 4780700 |
| Q12899;A2AE4 | Tripartite motif-containing protein 26 | TRIM26 | sp\|Q12899\|TRI26_HUMAN Tripartite motif-containing protein | 7 | 62,165 | 4,126 | | 2,001 | 3,0635 | 1519900 | 2525100 | 1427900 |
| Q99986;H0YJ5 | Serine/threonine-protein kinase VRK1 | VRK1 | sp\|Q99986\|VRK1_HUMAN Serine/threonine-protein kinase VR | 14 | 45,476 | 3,4612 | 3,0568 | 1,1048 | 3,0568 | 2738700 | 26700000 | 18471000 |
| C9J2Y9;P3087 | DNA-directed RNA polymerase;DNA-dir | POLR2B | tr\|C9J2Y9\|C9J2Y9_HUMAN DNA-directed RNA polymerase subu | 34 | 133,06 | 3,7497 | 3,0177 | 3,0513 | 3,0513 | 4282800 | 9848300 | 392790 |
| H3BV80;H3BM | RNA-binding protein with serine-rich d | RNPS1 | tr\|H3BV80\|H3BV80_HUMAN RNA-binding protein with serine- | 6 | 24,561 | 3,0441 | 3,3076 | 2,5543 | 3,0441 | 139510000 | 204410000 | 94293000 |
| Q9UBI6 | Guanine nucleotide-binding protein G( | GNG12 | sp\|Q9UBI6\|GBG12_HUMAN Guanine nucleotide-binding prote | 3 | 8,0061 | 3,0438 | 4,9367 | 2,0117 | 3,0438 | 3890300 | 34430000 | 84342000 |
| I1E4Y6;Q6Y7W | PERQ amino acid-rich with GYF domain | GIGYF2 | tr\|I1E4Y6\|I1E4Y6_HUMAN GRB10-interacting GYF protein 2 OS | 11 | 152,53 | 3,4506 | 2,6293 | | 3,03995 | 571610 | 1017200 | 0 |
| P16403;P1041 | Histone H1.2;Histone H1.4 | HIST1H1C;HIST1H | sp\|P16403\|H12_HUMAN Histone H1.2 OS=Homo sapiens OX=9 | 12 | 21,364 | 3,0264 | 2,2983 | 3,593 | 3,0264 | 103310000 | 303900000 | 129810000 |
| P19387;H3BR | DNA-directed RNA polymerase II subuni | POLR2C | sp\|P19387\|RPB3_HUMAN DNA-directed RNA polymerase II sub | 10 | 31,441 | 3,0233 | 3,8035 | 2,6379 | 3,0233 | 11207000 | 18173000 | 2879500 |
| Q5HYB6 | | DKFZp686J1372 | tr\|Q5HYB6\|Q5HYB6_HUMAN Epididymis luminal protein 189 ( | 21 | 27,175 | 4,5555 | 2,9635 | 2,9779 | 2,9779 | 1471400 | 6297900 | 1443000 |
| P06748;E5RI9 | Nucleophosmin | NPM1 | sp\|P06748\|NPM_HUMAN Nucleophosmin OS=Homo sapiens O | 17 | 32,575 | 3,5369 | 2,9192 | 1,7384 | 2,9192 | 45741000 | 124660000 | 102380000 |
| J3QLD9;E7EMK | Flotillin-2 | FLOT2 | tr\|J3QLD9\|J3QLD9_HUMAN Flotillin-2 OS=Homo sapiens OX=9 | 15 | 47,142 | 2,9067 | 4,1686 | 1,6969 | 2,9067 | 8440000 | 26271000 | 11510000 |
| P62879;C9JIS1 | Guanine nucleotide-binding protein G( | GNB2 | sp\|P62879\|GBB2_HUMAN Guanine nucleotide-binding protei | 12 | 37,331 | 2,9058 | 4,836 | 1,5201 | 2,9058 | 4881400 | 14551000 | 15388000 |
| Q53H12;E9PC1 | Acylglycerol kinase, mitochondrial | AGK | sp\|Q53H12\|AGK_HUMAN Acylglycerol kinase, mitochondrial C | 18 | 47,137 | 1,1258 | 2,8627 | 4,0535 | 2,8627 | 4820100 | 15152000 | 13972000 |
| Q13112 | Chromatin assembly factor 1 subunit B | CHAF1B | sp\|Q13112\|CAF1B_HUMAN Chromatin assembly factor 1 subu | 4 | 61,492 | 2,844 | 2,8809 | | 2,86245 | 512590 | 77757 | 0 |
| A0A087WVZ9; | DNA-directed RNA polymerases I, II, and | POLR2E | tr\|A0A087WVZ9\|A0A087WVZ9_HUMAN DNA-directed RNA po | 7 | 21,459 | 3,1167 | 2,4935 | | 2,8051 | 3322300 | 15849000 | 0 |
| Q92820 | Gamma-glutamyl hydrolase | GGH | sp\|Q92820\|GGH_HUMAN Gamma-glutamyl hydrolase OS=Hom | 8 | 35,964 | 2,8025 | 5,1253 | 2,0979 | 2,8025 | 2133200 | 7381800 | 1489700 |
| Q96GA3;A0A07 | Protein LTV1 homolog | LTV1 | sp\|Q96GA3\|LTV1_HUMAN Protein LTV1 homolog OS=Homo sa | 15 | 54,854 | 1,3874 | 3,0356 | 2,7805 | 2,7805 | 11848000 | 15414000 | 11793000 |
| P60709;A0A2R | Actin, cytoplasmic 1;Actin, cytoplasmi | ACTB | sp\|P60709\|ACTB_HUMAN Actin, cytoplasmic 1 OS=Homo sapie | 27 | 41,736 | 1,2178 | 3,5005 | 2,7751 | 2,7751 | 11323000 | 36019000 | 20782000 |
| Added01;CON__Q9U6Y5;I3L170 | | | tr\|Added01\|TauYfp_HUMAN Tau Yfp | 39 | 43,256 | 3,5063 | 2,7256 | 2,1606 | 2,7256 | 4130100000 | 8914000000 | 3833400000 |
| Q99613;B5ME1 | Eukaryotic translation initiation factor | EIF3C;EIF3CL | sp\|Q99613\|EIF3C_HUMAN Eukaryotic translation initiation fac | 33 | 105,34 | 2,5577 | 2,8357 | | 2,6967 | 855670 | 4670000 | 443520 |
| P28066 | Proteasome subunit alpha type-5 | PSMA5 | sp\|P28066\|PSA5_HUMAN Proteasome subunit alpha type-5 OS | 13 | 26,411 | | 3,108 | 2,254 | 2,681 | 713200 | 3627100 | 526020 |
| Q9HCM4 | Band 4.1-like protein 5 | EPB41L5 | sp\|Q9HCM4\|E41L5_HUMAN Band 4.1-like protein 5 OS=Homo | 9 | 81,855 | 2,1785 | 3,0182 | | 2,59835 | 227100 | 3722200 | 0 |
| A0A087WYV5; | Slit homolog 2 protein;Slit homolog 2 | SLIT2 | tr\|A0A087WYV5\|A0A087WYV5_HUMAN Slit homolog 2 protei | 9 | 159,98 | 2,0219 | 3,1071 | | 2,5645 | 487410 | 1400300 | 599630 |
| A0A1W2PQ90; | DNA repair protein RAD50 | RAD50 | tr\|A0A1W2PQ90\|A0A1W2PQ90_HUMAN Uncharacterized pro | 25 | 142,94 | 2,5131 | 3,214 | 0,89296 | 2,5131 | 1048700 | 5250600 | 1514700 |
| Q9H0U4;E9PLD | Ras-related protein Rab-1B;Putative Ra | RAB1B;RAB1C | sp\|Q9H0U4\|RAB1B_HUMAN Ras-related protein Rab-1B OS=Ho | 12 | 22,171 | | 2,7446 | 2,1603 | 2,45245 | 682330 | 3902100 | 564680 |
| Q13823;H0YG1 | Nucleolar GTP-binding protein 2 | GNL2 | sp\|Q13823\|NOG2_HUMAN Nucleolar GTP-binding protein 2 O | 18 | 83,654 | 2,5228 | 2,3703 | | 2,44655 | 504360 | 3938100 | 57885 |
| Q07021;I3L3Q | Complement component 1 Q subcomp | C1QBP | sp\|Q07021\|C1QBP_HUMAN Complement component 1 Q subc | 10 | 31,362 | 2,1285 | 5,1114 | 2,4395 | 2,4395 | 5133600 | 39902000 | 12700000 |
| Q7Z417 | Nuclear fragile X mental retardation-int | NUFIP2 | sp\|Q7Z417\|NUFP2_HUMAN Nuclear fragile X mental retardatic | 15 | 76,12 | 2,9962 | 2,4364 | 2,0328 | 2,4364 | 5829600 | 7837800 | 5467700 |
| P08670;B0YJC | Vimentin | VIM | sp\|P08670\|VIME_HUMAN Vimentin OS=Homo sapiens OX=960 | 45 | 53,651 | 2,4027 | 2,6728 | 1,6894 | 2,4027 | 59024000 | 131930000 | 146780000 |
| P19525;C9JZT | Interferon-induced, double-stranded R | EIF2AK2 | sp\|P19525\|E2AK2_HUMAN Interferon-induced, double-strand | 12 | 62,094 | 2,5648 | 2,384 | 2,0421 | 2,384 | 145870 | 749090 | 114650 |
| Q01082 | Spectrin beta chain, non-erythrocytic 1 | SPTBN1 | sp\|Q01082\|SPTB2_HUMAN Spectrin beta chain, non-erythrocy | 120 | 274,61 | 2,354 | 5,0002 | 1,5658 | 2,354 | 14363000 | 46226000 | 27935000 |
| A0A0D9SF54;A0A0D9SGF6;A0A0D9SFF6;A0A0D9SFH4; | | SPTAN1 | tr\|A0A0D9SF54\|A0A0D9SF54_HUMAN Spectrin alpha chain, n | 132 | 282,83 | 2,3502 | 5,0256 | 1,4746 | 2,3502 | 16833000 | 62862000 | 26440000 |
| P07948;E5RJ3 | Tyrosine-protein kinase Lyn | LYN | sp\|P07948\|LYN_HUMAN Tyrosine-protein kinase Lyn OS=Hom | 12 | 58,573 | | 2,3857 | 2,2951 | 2,3404 | 0 | 1032300 | 2217900 |
| Q9UN86;D6RA | Ras GTPase-activating protein-binding | G3BP2 | sp\|Q9UN86\|G3BP2_HUMAN Ras GTPase-activating protein-bin | 11 | 54,12 | 2,3338 | 1,3159 | 2,5495 | 2,3338 | 8272900 | 13070000 | 6033900 |
| P07900;G3V2J | Heat shock protein HSP 90-alpha | HSP90AA1 | sp\|P07900\|HS90A_HUMAN Heat shock protein HSP 90-alpha C | 63 | 84,659 | 2,277 | 4,0858 | 2,008 | 2,277 | 6729400 | 12647000 | 4596800 |
| P46013 | Antigen KI-67 | MKI67 | sp\|P46013\|KI67_HUMAN Proliferation marker protein Ki-67 O | 17 | 358,69 | 2,3191 | 2,2468 | 2,2169 | 2,2468 | 524560 | 475360 | 44507 |

| | | | | | | | | | | | | |
|---|---|---|---|---|---|---|---|---|---|---|---|---|
| P60228;E5RGA | Eukaryotic translation initiation factor | EIF3E | sp\|P60228\|EIF3E_HUMAN Eukaryotic translation initiation fac | 25 | 52,22 | 2,3804 | 2,2127 | 1,9171 | 2,2127 | 1089300 | 1573400 | 1367200 |
| P49916;K7ERZ! | DNA ligase 3 | LIG3 | sp\|P49916\|DNLI3_HUMAN DNA ligase 3 OS=Homo sapiens OX= | 8 | 112,91 | 2,9178 | 2,203 | 2,1767 | 2,203 | 1532800 | 1039000 | 946440 |
| Q5RKV6 | Exosome complex component MTR3 | EXOSC6 | sp\|Q5RKV6\|EXOS6_HUMAN Exosome complex component MTI | 6 | 28,235 | 2,3573 | 2,1818 | 2,111 | 2,1818 | 11071000 | 15292000 | 4620400 |
| P12931 | Proto-oncogene tyrosine-protein kinas | SRC | sp\|P12931\|SRC_HUMAN Proto-oncogene tyrosine-protein kin | 8 | 59,834 | 2,168 | 4,1415 | 1,8886 | 2,168 | 748320 | 2601200 | 1702500 |
| P63092;Q5JWI | Guanine nucleotide-binding protein G( | GNAS | sp\|P63092\|GNAS2_HUMAN Guanine nucleotide-binding prote | 13 | 45,664 | 2,1627 | 4,1417 | 1,5402 | 2,1627 | 1488700 | 9048500 | 3740200 |
| O75531;E9PJJ8 | Barrier-to-autointegration factor;Barri | BANF1 | sp\|O75531\|BAF_HUMAN Barrier-to-autointegration factor OS= | 6 | 10,058 | 2,1605 | 2,4895 | 1,4442 | 2,1605 | 167700000 | 464450000 | 175040000 |
| Q9Y265;E7ETR( | RuvB-like 1 | RUVBL1 | sp\|Q9Y265\|RUVB1_HUMAN RuvB-like 1 OS=Homo sapiens OX= | 24 | 50,227 | 1,8486 | 2,1597 | 2,5745 | 2,1597 | 2800700 | 6342500 | 382330 |
| O14578;H7BYJ | Citron Rho-interacting kinase | CIT | sp\|O14578\|CTRO_HUMAN Citron Rho-interacting kinase OS=H | 11 | 231,43 | 3,1776 | 2,1397 | 0,96079 | 2,1397 | 57864 | 292140 | 49921 |
| P27986;H0YBC | Phosphatidylinositol 3-kinase regulato | PIK3R1 | sp\|P27986\|P85A_HUMAN Phosphatidylinositol 3-kinase regul | 6 | 83,597 | 1,5657 | 2,1169 | 2,5085 | 2,1169 | 745890 | 1116000 | 1210500 |
| Q6WCQ1;J3KS\ | Myosin phosphatase Rho-interacting pr | MPRIP | sp\|Q6WCQ1\|MPRIP_HUMAN Myosin phosphatase Rho-interac | 12 | 116,53 | 2,1049 | 2,173 | 0,79734 | 2,1049 | 588250 | 1624600 | 104110 |
| P08754 | Guanine nucleotide-binding protein G( | GNAI3 | sp\|P08754\|GNAI3_HUMAN Guanine nucleotide-binding prote | 15 | 40,532 | 2,0157 | 4,9059 | 2,0813 | 2,0813 | 19273000 | 75469000 | 42160000 |
| O15234;J3KSY | Protein CASC3 | CASC3 | sp\|O15234\|CASC3_HUMAN Protein CASC3 OS=Homo sapiens C | 8 | 76,277 | 2,067 | 2,0138 | 2,7423 | 2,067 | 698630 | 2073400 | 4126900 |
| P17987;E7EQR | T-complex protein 1 subunit alpha | TCP1 | sp\|P17987\|TCPA_HUMAN T-complex protein 1 subunit alpha C | 34 | 60,343 | 1,5294 | 2,2873 | 2,0582 | 2,0582 | 485300 | 1703400 | 621710 |
| Q5SRQ6;P6787 | Casein kinase II subunit beta | CSNK2B;CSNK2B- | tr\|Q5SRQ6\|Q5SRQ6_HUMAN Casein kinase II subunit beta OS=H | 8 | 26,925 | 2,0436 | 2,4349 | 1,2233 | 2,0436 | 5899500 | 15187000 | 987590 |
| P08238;Q58FF | Heat shock protein HSP 90-beta | HSP90AB1 | sp\|P08238\|HS90B_HUMAN Heat shock protein HSP 90-beta OS | 64 | 83,263 | 1,9891 | 2,9312 | 2,02 | 2,02 | 7615400 | 29148000 | 15600000 |
| O95425;A0A0J | Supervillin | SVIL | sp\|O95425\|SVIL_HUMAN Supervillin OS=Homo sapiens OX=96 | 9 | 247,74 | 1,4281 | 3,0818 | 2,0134 | 2,0134 | 360680 | 1636100 | 213390 |
| Q14676;A2AB0 | Mediator of DNA damage checkpoint pr | MDC1 | sp\|Q14676\|MDC1_HUMAN Mediator of DNA damage checkpoi | 20 | 226,66 | 2,1653 | 2,0075 | 1,5305 | 2,0075 | 303620 | 152960 | 60101 |
| Q14008;H0YDX | Cytoskeleton-associated protein 5 | CKAP5 | sp\|Q14008\|CKAP5_HUMAN Cytoskeleton-associated protein 5 | 72 | 225,49 | 1,7242 | 2,0021 | 2,3599 | 2,0021 | 325010 | 2252600 | 1461800 |
